# The relative coding strength of object identity and nonidentity features in human occipito-temporal cortex and convolutional neural networks

**DOI:** 10.1101/2020.08.11.246967

**Authors:** Yaoda Xu, Maryam Vaziri-Pashkam

## Abstract

Any given visual object input is characterized by multiple visual features, such as identity, position and size. Despite the usefulness of identity and nonidentity features in vision and their joint coding throughout the primate ventral visual processing pathway, they have so far been studied relatively independently. Here we document the relative coding strength of object identity and nonidentity features in a brain region and how this may change across the human ventral visual pathway. We examined a total of four nonidentity features, including two Euclidean features (position and size) and two non-Euclidean features (image statistics and spatial frequency content of an image). Overall, identity representation increased and nonidentity feature representation decreased along the ventral visual pathway, with identity outweighed the non-Euclidean features, but not the Euclidean ones, in higher levels of visual processing. A similar analysis was performed in 14 convolutional neural networks (CNNs) pretrained to perform object categorization with varying architecture, depth, and with/without recurrent processing. While the relative coding strength of object identity and nonidentity features in lower CNN layers matched well with that in early human visual areas, the match between higher CNN layers and higher human visual regions were limited. Similar results were obtained regardless of whether a CNN was trained with real-world or stylized object images that emphasized shape representation. Together, by measuring the relative coding strength of object identity and nonidentity features, our approach provided a new tool to characterize feature coding in the human brain and the correspondence between the brain and CNNs.

**SIGNIFICANCE STATEMENT:** This study documented the relative coding strength of object identity compared to four types of nonidentity features along the human ventral visual processing pathway and compared brain responses with those of 14 CNNs pretrained to perform object categorization. Overall, identity representation increased and nonidentity feature representation decreased along the ventral visual pathway, with the coding strength of the different nonidentity features differed at higher levels of visual processing. While feature coding in lower CNN layers matched well with that of early human visual areas, the match between higher CNN layers and higher human visual regions were limited. Our approach provided a new tool to characterize feature coding in the human brain and the correspondence between the brain and CNNs.

## INTRODUCTION

In real-world vision, object identity information always appears together with nonidentity information, such as the position and size of an object. Nevertheless, vision research in the past several decades has mainly focused on discarding nonidentity features to recover object identity representations regardless of how objects may appear in the real world. The formation of such identity-preserving transformation tolerant object representations has been regarded as the defining feature of primate high-level vision (DiCarlo & Cox, 2007; DiCarlo et al., 2012).

Vision, however, is not just about object recognition; it also helps us to interact with the objects in the world. For example, to pick up an object, its position, size and orientation, rather than its identity, would be most relevant. In fact, both object identity and nonidentity features are represented together in higher visual areas in occipito-temporal cortex (OTC) in both macaques (Hung et al., 2005; Hong et al., 2016) and humans (Schwarzlose et al., 2008; Kravitz et al., 2008 & 2010; Carlson et al., 2011; Cichy et al., 2011 & 2013; Vaziri-Pashkam et al., 2019), with these two types of information being represented in a largely distributed and overlapping manner (Hong et al., 2016). Despite the usefulness of object identity and nonidentity features in vision and their joint coding in visual processing, they have so far been studied relatively independently. What is the relative coding strength of these two types of information within a brain region? How does it change over the course of visual processing in different brain regions? Answers to these questions will not only better characterize the coding of different visual features within a brain region, but also delineate the evolution of visual information representation along the ventral visual processing pathway.

Recent hierarchical CNNs have achieved human-like object categorization performance (Kriegeskorte, 2015; Yamins & Dicarlo, 2016; Rajalingham, et al., 2018; Serre, 2019), with representations formed in early and late layers of the network tracking those of the human early and later visual processing regions, respectively (Khaligh-Razavi & Kriegeskorte, 2014; Güçlü & van Gerven, 2015; Cichy et al., 2016; Eickenberg et al., 2017). Together with results from monkey neurophysiological studies, CNNs have been regarded by some as the current best models of the primate visual system (e.g., Cichy & Kaiser, 2019; Kubilius et al., 2019). Nevertheless, we lack a detailed and clear understanding of how information is processed in CNNs. This is especially evident from recent studies reporting a number of discrepancies in processing between the brain and CNNs (Serre, 2019). Our own investigation has shown that the close brain-CNN correspondence was rather limited and could not fully capture visual processing in the human brain (Xu & Vaziri-Pashkam, 2020). By examining the representations of object identity and nonidentity features in CNNs and comparing the results with those from the human brain, we can form a deeper understanding of how visual information is processed in CNNs.

In this study we analyzed four existing fMRI data sets and documented the relative encoding strength of object identity and nonidentity features and its evolution across the human ventral visual processing pathway in OTC regions (Figure 1). We compared object identity representation with four types of nonidentity features, including two Euclidean features (position and size) and two non-Euclidean features (image statistics and the spatial frequency (SF) content of an image). We found an overall increase of identity and a decrease of nonidentity information representation along the human visual processing hierarchy. While identity representation dominated those of the non-Euclidean features in higher levels of visual processing, this was not the case for the Euclidean features examined. We additionally examined 14 different CNNs pretrained to perform object categorization with varying architecture, depth, and with/without recurrent processing. We found that while the relative coding strength of object identity and nonidentity features in lower CNN layers matched with that in human early visual areas, the match between higher CNN layers and higher human visual regions were limited.

**Figure 1.**
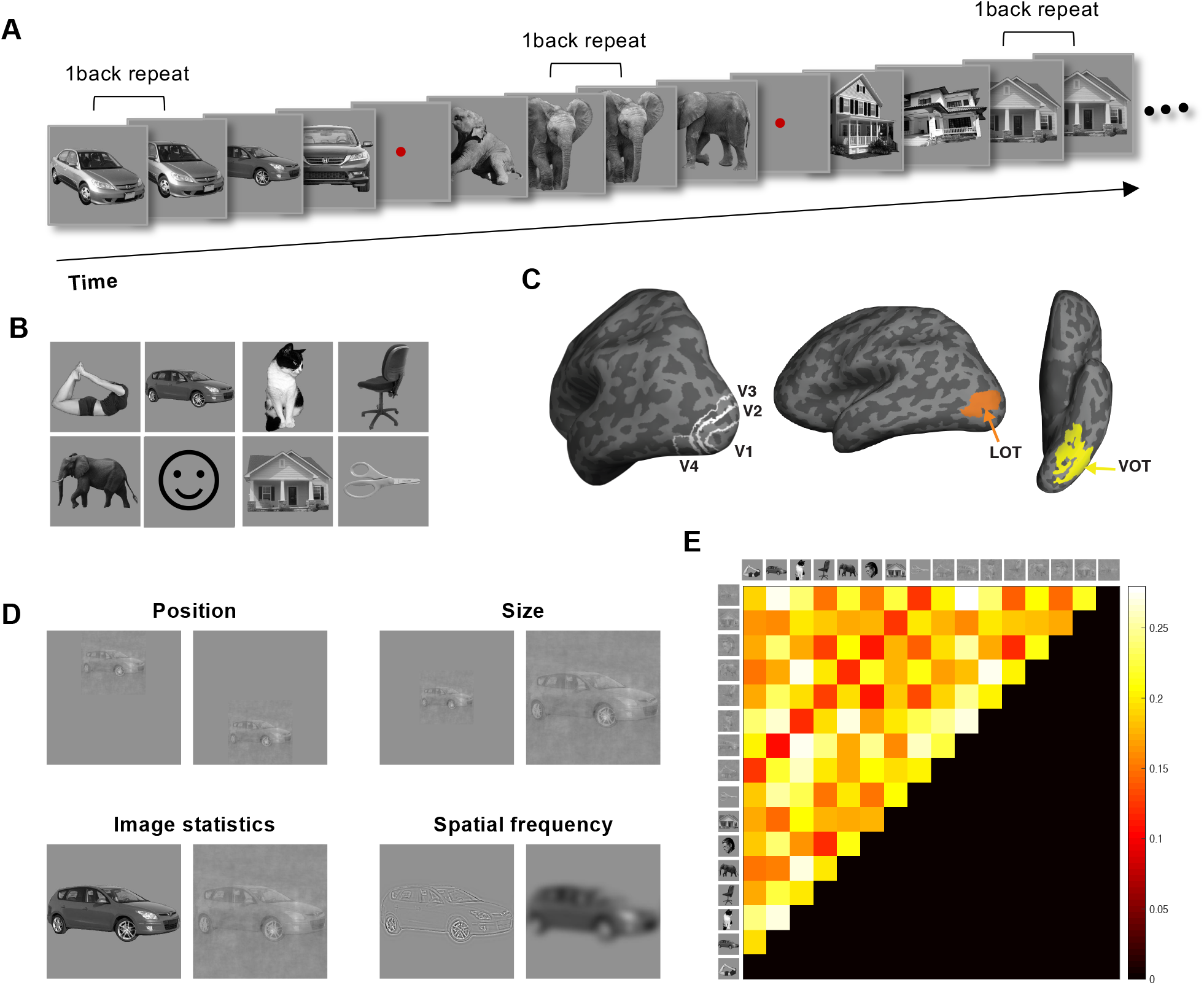
**(A)** An illustration of the block design paradigm used. Participants performed a one-back repetition detection task on the images. An actual block in the experiment contained 10 images with two repetitions per block. See Methods for more details. **(B)** The eight real-world object categories used. **(C)** The brain regions examined. They included topographically defined early visual areas V1 to V4 and functionally defined higher object processing regions LOT and VOT. **(D)** The four types of nonidentity transformations examined. They included two Euclidean transformations - position and size, and two non-Euclidean transformations - image stats and spatial frequency. **(E)** An example RDM from a brain region containing all pairwise Euclidean distances for all the categories shown across both values of the image stats feature (i.e., original and controlled).

## Materials and Methods

### fMRI Experimental Details

Details of the fMRI experiments have been described in two previously published studies (Vaziri-Pashkam & Xu, 2019 and Vaziri-Pashkam et al., 2019). They are summarized here for the readers’ convenience.

Seven, seven, six, and ten healthy human participants with normal or corrected to normal visual acuity, all right-handed, and aged between 18-35 took part in Experiments 1 to 4, respectively. Each main experiment was performed in a separate session lasting between 1.5 and 2 hours. Each participant also completed two additional sessions for topographic mapping and functional localizers. MRI data were collected using a Siemens MAGNETOM Trio, A Tim System 3T scanner, with a 32-channel receiver array head coil. For all the fMRI scans, a T2*-weighted gradient echo pulse sequence with TR of 2 sec and voxel size of 3 mm × 3 mm × 3 mm was used. FMRI data were analyzed using FreeSurfer (surfer.nmr.mgh.harvard.edu), FsFast (Dale et al., 1999) and in-house MATLAB codes. FMRI data preprocessing included 3D motion correction, slice timing correction and linear and quadratic trend removal. Following standard practice, a general linear model was then applied to the fMRI data to extract beta weights as response estimates.

The general experimental paradigm consisted of a 1-back image repetition detection task in which participants viewed a stream of sequentially presented images and pressed a response button when the same image repeated back to back (Figure 1a). This task engaged participants’ attention on the object shapes and ensured robust fMRI responses. Two image repetitions occurred randomly in each image block. We used cut-out grey-scaled images from eight real-world object categories (faces, bodies, houses, cats, elephants, cars, chairs, and scissors) and modified them to occupy roughly the same area on the screen (Figure 1b). For each object category, we selected ten exemplar images that varied in identity, pose and viewing angle to minimize the low-level similarities among them. Each block of image presentation contained images from the same object category. Participants fixated at a central red dot throughout the experiment. Eye-movements were monitored in all the fMRI experiments to ensure proper fixation. We examined responses from early visual areas V1 to V4 and higher visual processing regions in lateral occipito-temporal (LOT) and ventral occipito-temporal (VOT) cortex (see more details below) (Figure 1c).

In Experiment 1, we tested position tolerance and presented images either above or below the fixation (Figure 1d). Each block contained a random sequential presentation of ten exemplars from the same object category shown either all above or all below the fixation. To ensure that object identity representation in lower brain regions truly reflected the representation of object identity and not low-level differences among the images of the different categories, controlled images with low-level image differences equated among the different categories were shown. These controlled images were generated by equalizing contrast, luminance and spatial frequency of the images across all the categories using the shine toolbox (Willenbockel et al., 2010, see Figure 1c). All images subtended 2.9° × 2.9° and were shown at 1.56° above the fixation in half of the 16 blocks and the same distance below the fixation in the other half of the blocks. Each image was presented for 200 msec followed by a 600 msec blank interval between the images. Each experimental run contained 16 blocks, one for each of the 8 categories in each of the two image positions. The order of the eight object categories and the two positions were counterbalanced across runs and participants. Each block lasted 8 secs and followed by an 8-sec fixation period. There was an additional 8-sec fixation period at the beginning of the run. Each participant completed one scan session with 16 runs for this experiment, each lasting 4 mins 24 secs.

In Experiment 2, we tested size tolerance and presented images either in a large size (5.77° × 5.77°) or small size (2.31° × 2.31°) centered at fixation (Figure 1d). As in Experiment 1, controlled images were used here. Half of the 16 blocks contained small images and the other half, large images. Other details of the experiment were identical to that of Experiment 1.

In Experiment 3, we tested image stats tolerance and presented images at fixation either in the original unaltered format or in the controlled format (subtended 4.6° × 4.6°) (Figure 1d). Half of the 16 blocks contained original images and the other half, controlled images. Other details of the experiment were identical to that of Experiment 1.

In Experiment 4, only six of the original eight object categories were included and they were faces, bodies, houses, elephants, cars, and chairs. Images were shown in 3 conditions: Full-SF, High-SF, and Low-SF (Figure 1d). In the Full-SF condition, the full spectrum images were shown without modification of the SF content. In the High-SF condition, images were high-pass filtered using an FIR filter with a cutoff frequency of 4.40 cycles per degree. In the Low-SF condition, the images were low-pass filtered using an FIR filter with a cutoff frequency of 0.62 cycles per degree. The DC component was restored after filtering so that the image backgrounds were equal in luminance. Each run contained 18 blocks, one for each of the category and SF condition combination. Each participant completed a single scan session containing 18 experimental runs, each lasting 5 minutes. Other details of the experimental design were identical to that of Experiment 1. Only the results from the LF and HF conditions were included in the present analysis.

We examined responses from independent localized early visual areas V1 to V4 and higher visual processing regions LOT and VOT (Figure 1c). V1 to V4 were mapped with flashing checkerboards using standard techniques (Sereno et al., 1995). Following the detailed procedures described in Swisher et al. (2007) and by examining phase reversals in the polar angle maps, we identified areas V1 to V4 in the occipital cortex of each participant (see also Bettencourt & Xu, 2016) (Figure 1c). To identify LOT and VOT, following Kourtzi and Kanwisher (2000), participants viewed blocks of face, scene, object and scrambled object images. These two regions were then defined as a cluster of continuous voxels in the lateral and ventral occipital cortex, respectively, that responded more to the original than to the scrambled object images. LOT and VOT loosely correspond to the location of LO and pFs (Malach et al., 1995; Grill-Spector et al.,1998; Kourtzi & Kanwisher, 2000) but extend further into the temporal cortex in an effort to include as many object-selective voxels as possible in occipito-temporal regions.

To generate the fMRI response pattern for each ROI in a given run, we first convolved an 8-second stimulus presentation boxcar (corresponding to the length of each image block) with a hemodynamic response function to each condition; we then conducted a general linear model analysis to extract the beta weight for each condition in each voxel of that ROI. These voxel beta weights were used as the fMRI response pattern for that condition in that run. Following Tarhan and Konkle (2019), to increase power, we selected the top 75 most reliable voxels in each ROI for further analyses. This was done by splitting the data into odd and even halves, averaging the data across the runs within each half, correlating the beta weights from all the conditions between the two halves for each voxel, and then selecting the top 75 voxels showing the highest correlation. This is akin to including the best units in monkey neurophysiological studies. For example, Cadieu et al. (2014) only selected a small subset of all recorded single units for their brain-CNN analysis. We obtained the fMRI response pattern for each condition from the 75 most reliable voxels in each ROI of each run. We then averaged the fMRI response patterns from all the runs and applied z-normalization to the averaged pattern for each condition in each ROI to remove amplitude differences between conditions and ROIs before further analyses were carried out. Very similar results were obtained if we included all voxels in an ROI instead of just the 75 most reliable voxels (see Extended Figure 3-1).

### CNN details

We included 14 CNNs in our analyses (see Table 1). They included both shallower networks, such as Alexnet, VGG16 and VGG 19, and deeper networks, such as Googlenet, Inception-v3, Resnet-50 and Resnet-101. We also included a recurrent network, Cornet-S, that has been shown to capture the recurrent processing in macaque IT cortex with a shallower structure (Kubilius et al., 2019; Kar et al., 2019). This CNN has been recently argued to be the current best model of the primate ventral visual processing regions (Kar et al., 2019). All the CNNs used were trained with ImageNet images (Deng et al., 2009).

**Table 1.** The CNNs and the layers examined in this study.

| CNN name | Depth/Blocks | Layers | N of Layers Sampled | Sampled Layer Names and Locations (indicated in the parenthesis) |
| --- | --- | --- | --- | --- |
| Alexnet | 8 | 25 | 6 | 'pool1' (5), 'pool2' (9), 'pool5' (16), 'fc6' (17), 'fc7' (20), 'fc8' (23) |
| Cornet-S | 4 | 42 | 6 | 'V1_output' (8), 'V2_output' (18), 'V4_output' (28), 'IT_output' (38), 'decoder_avgpool' (39), 'decoder_output' (42) |
| Densenet-201 | 201 | 709 | 6 | 'pool1' (6), 'pool2_pool' (52), 'pool3_pool' (140), 'pool4_pool' (480), 'avg_pool' (706), 'fc1000' (707) |
| Googlenet | 22 | 144 | 6 | 'pool1-3x3_s2' (4), 'pool2-3x3_s2' (11), 'pool3-3x3_s2' (40), 'pool4-3x3_s2' (111), 'pool5-7x7_s1' (140), 'loss3-classifier' (142) |
| Inception_v3 | 48 | 316 | 11 | 'average_pooling2d_1' (29), 'average_pooling2d_2' (52), 'average_pooling2d_3' (75), 'average_pooling2d_4' (121), 'average_pooling2d_5' (153), 'average_pooling2d_6' (185), 'average_pooling2d_7' (217), 'average_pooling2d_8' (264), 'average_pooling2d_9' (295), 'avg_pool' (313), 'predictions' (314) |
| Inception-resnet_v2 | 164 | 825 | 7 | 'max_pooling2d_1' (12), 'max_pooling2d_2' (19), 'average_pooling2d_1' (29), 'max_pooling2d_3' (285), 'max_pooling2d_4' (648), 'avg_pool' (822), 'predictions' (823) |
| Mobilenet_v2 | 54 | 155 | 10 | 'block_2_project_BN' (26), 'block_4_project_BN' (43), 'block_6_project_BN' (61), 'block_8_project_BN' (78), 'block_10_project_BN' (96), 'block_12_project_BN' (113), 'block_14_project_BN' (130), 'block_16_project_BN' (148), 'global_average_pooling2d_1' (152), 'Logits' (153) |
| Resnet-18 | 18 | 72 | 6 | 'pool1' (6), 'res2b_relu' (20), 'res3b_relu' (36), 'res4b_relu' (52), 'pool5' (69), 'fc1000' (70) |
| Resnet-50 | 50 | 177 | 6 | 'max_pooling2d_1' (5), 'activation_10_relu' (37), 'activation_22_relu' (79), 'activation_40_relu' (141), 'avg_pool' (174), 'fc1000' (175) |
| Resnet-101 | 101 | 347 | 6 | 'pool1' (5), 'res2c_relu' (37), 'res3b3_relu' (79), 'res4b22_relu' (311), 'pool5' (344), 'fc1000' (345) |
| Squeezenet | 18 | 68 | 5 | 'pool1' (4), 'pool3' (19), 'pool5' (34), 'conv10' (64), 'pool10' (66) |
| Vgg-16 | 16 | 41 | 8 | 'pool1' (6), 'pool2' (11), 'pool3' (18), 'pool4' (25), 'pool5' (32), 'fc6' (33), 'fc7' (36), 'fc8' (39) |
| Vgg-19 | 19 | 47 | 8 | 'pool1' (6), 'pool2' (11), 'pool3' (20), 'pool4' (29), 'pool5' (38), 'fc6' (39), 'fc7' (42), 'fc8' (45) |
| Xception | 71 | 171 | 8 | 'block2_pool' (18), 'block4_pool' (42), 'block6_sepconv3_bn' (68), 'block8_sepconv3_bn' (94), 'block10_sepconv3_bn' (120), 'block12_sepconv3_bn' (146), 'avg_pool' (168), 'predictions' (169) |

To understand how the specific training images would impact CNN representations, besides CNNs trained with ImageNet images, we also examined Resnet-50 trained with stylized ImageNet images (Geirhos et al., 2019). We examined the representations formed in Resnet-50 pretrained with three different procedures (Geirhos et al., 2019): trained only with the stylized ImageNet Images (RN50-SIN), trained with both the original and the stylized ImageNet Images (RN50-SININ), and trained with both sets of images and then fine-tuned with the stylized ImageNet images (RN50-SININ-IN).

Following O’Connor et al. (2018), we sampled between 6 and 11 mostly pooling and FC layers of each CNN (see Supplemental Table 1 for the specific CNN layers sampled). Pooling layers were selected because they typically mark the end of processing for a block of layers before information is pooled and passed on to the next block of layers. When there were no obvious pooling layers present, the last layer of a block was chosen. For a given CNN layer, we extracted the CNN layer output for each object image in a given condition, averaged the output from all images in a given category for that condition, and then z-normalized the responses to generate the CNN layer response for that object category in that condition (similar to how fMRI category responses were extracted). Cornet-S and the different versions of Resnet-50 were implemented in Python. All other CNNs were implemented in Matlab. Output from all CNNs were analyzed and compared with brain responses using Matlab.

### Visualizing and quantifying the relative coding strength of identity and nonidentity features

To directly visualize how object identity and nonidentity features may be represented together in a brain region, from the z-normalized fMRI response patterns averaged over all the runs, we first calculated all pairwise Euclidean distances including all the categories shown and both values of each nonidentity feature (e.g., small and large). We then constructed a category representational dissimilarity matrix (RDM) for each brain region (Kriegeskorte & Kievit, 2013, see Figure 1e). Using multi-dimensional scaling (MDS, Shepard, 1980), we visualized this RDM by projecting the first two dimensions that captured most of the variance onto a 2D space with the distance denoting the similarity in representation among the different feature combinations (Figure 2 and Extended Figures 2-1 to 2-4).

**Figure 2.**
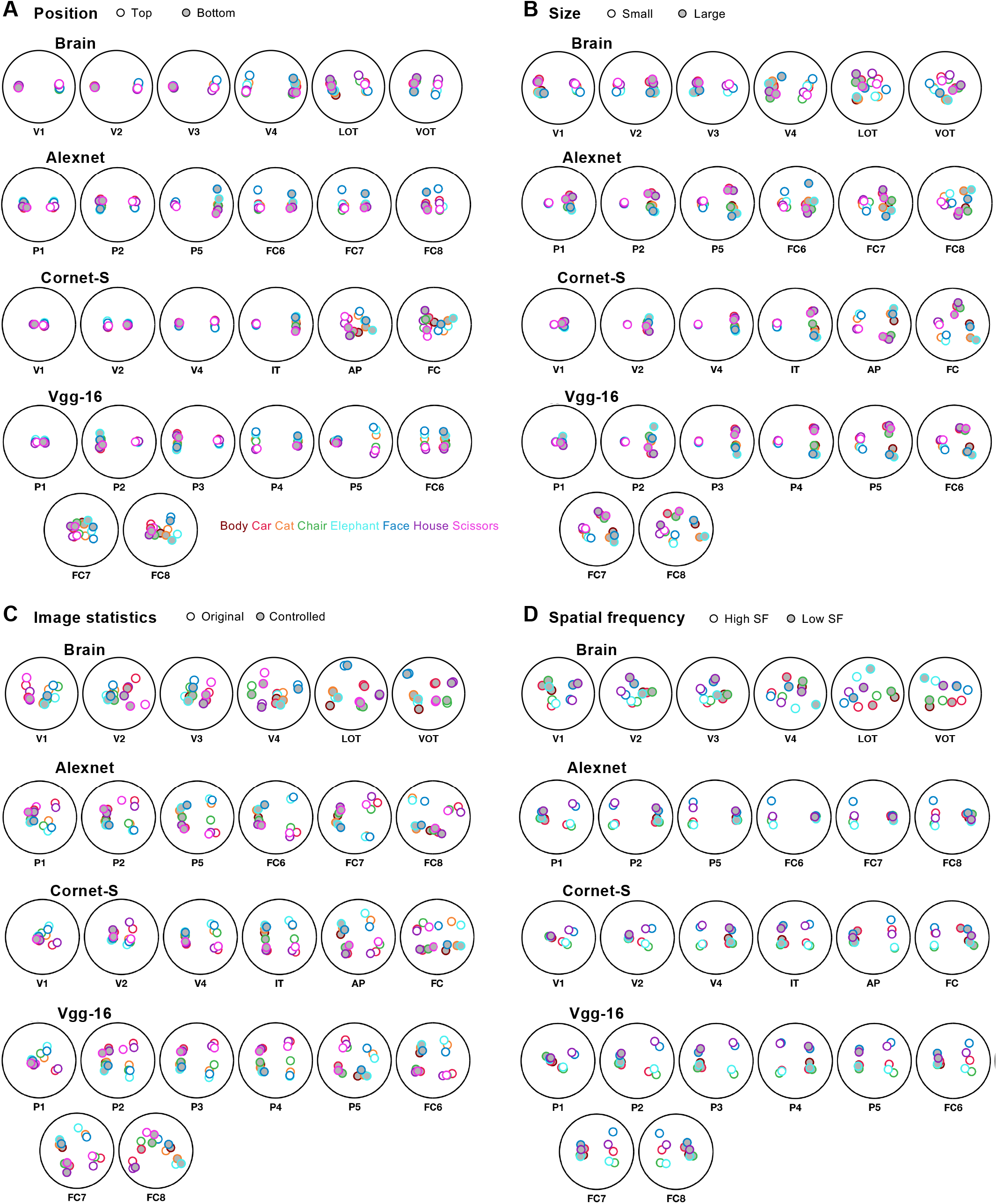
Visualizing feature coding in human OTC and CNNs. Using MDS, the RDMs containing object categories in both values of a nonidentity feature from each brain region/sampled CNN layer was visualized. This was done by projecting the first two dimensions that captured most of the variance onto a 2D space with the distance denoting the similarity in representation among the different feature combinations.

To quantify the relative coding strength of the different features in a brain region, from the z-normalized fMRI response patterns, we obtained the averaged Euclidean distance between different object categories sharing the same value of a nonidentity feature (*d_within_*) and the averaged Euclidean distance between the same object category across the two values of a nonidentity feature (*d_between_*). We then constructed an *identity dominance index* (*ODI*) as:

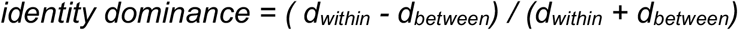

An identity-only representation that disregards the nonidentity feature would have a *d_between_* of “0” and an *identity dominance* of “1”. Conversely, a nonidentity-only representation that disregards identity would have a *d_within_* of “0” and an *identity dominance* of “-1”. An *identity dominance* of “0” indicates that equal representational strength of object identity and nonidentity features, such that an object category is equally similar to itself in the other value of the nonidentity feature as it is to the other categories sharing the same value of the nonidentity feature. Figures 3a and 3b illustrate two scenarios of how the relative coding strength of object identity and nonidentity features may change *identity dominance* between two hypothetical brain regions. A similar procedure was applied to CNN layer output to calculate *identity dominance* in CNN outputs.

**Figure 3.**
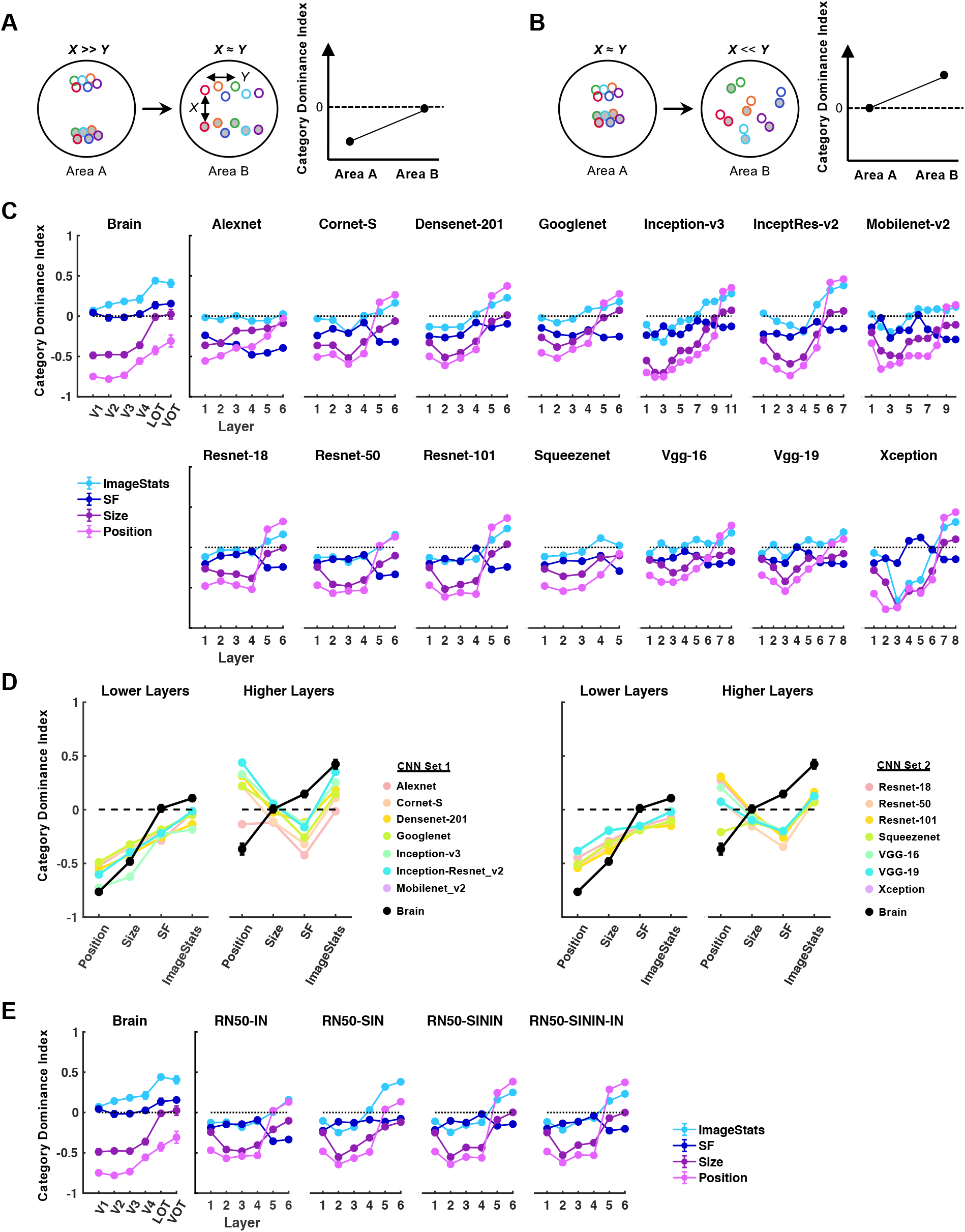
Quantifying the relative coding strength of identity and nonidentity features in human OTC and CNNs. An *identity dominance* index was used to quantify the relative coding strength of object identity and nonidentity features in a brain region and a CNN layer (see Methods). An identity-only representation that disregards the nonidentity feature would an *identity dominance* of “1”. Conversely, a nonidentity-only representation that disregards identity would have an *identity dominance* of “-1”. **(A)** and **(B)** Two scenarios of how a change in the relative coding strength of identity and nonidentity features in two hypothetical brain regions may change the resulting *identity dominance* measure. **(C)***Identity dominance* measures from human OTC and CNNs comparing the coding of identify with each of the four nonidentity features. **(D)** Direct comparisons of *identity dominance* across the four types of nonidentity features. For brain regions, only the average of the two early brain regions (V1 and V2) and the two higher brain regions (LOT and VOT) were included. For CNNs, only the average of the first two sampled layers and the average of the last two sampled layers were included. **(E)***Identity dominance* from human OTC and Resnet-50 trained with original and stylized ImageNet images. Resnet-50 was pretrained either with the original ImageNet images (RN50-IN), the stylized ImageNet Images (RN50-SIN), both the original and the stylized ImageNet Images (RN50-SININ), or both sets of images and then fine-tuned with the stylized ImageNet images (RN50-SININ-IN).

All reported t tests were two-tailed and corrected for multiple comparisons using the Benjamini and Hochberg method (Benjamini & Hochberg, 1995).

## RESULTS

Any given visual object input is characterized by multiple visual features, such as its identity, position and size. Although object identity representation has been intensely studied in the primate brain, nonidentity features, such as position and size, have also been found to be robustly coded throughout the primate ventral visual processing pathway (Hong et al., 2016; Vaziri-Pashkam et al., 2019). Here we document the relative coding strength of object identity and nonidentity features in a brain region and how this may change across the ventral visual processing pathways in human OTC. We also examined responses from 14 CNNs pretrained to perform object categorization with varying architecture, depth and the presence/absence of recurrent processing. Our fMRI data were collected with a block design in which responses were averaged over a whole block of multiple exemplars from the same category to increase SNR (Figure 1a). A total of eight real-world object categories were included and they were bodies, cars, cats, chairs, elephants, faces, houses, and scissors (Vaziri-Pashkam & Xu, 2019; Vaziri-Pashkam et al., 2019; Figure 1b). The images were shown in conjunction with four nonidentity features (Figure 1d), including two Euclidean features - position (top vs bottom), size (small vs large), and two non-Euclidean features image - statistics (original vs controlled) and SF of an image (high SF vs low SF). Controlled images were created to achieve spectrum, histogram, and intensity normalization and equalization across the images from the different categories (Willenbockel et al., 2010). Controlled images also appeared in the position and size manipulations to ensure that object identity representation in lower brain regions would reflect the representation of object identity and not low-level differences among the images of the different categories.

To increase power, we extracted the averaged neural response patterns from each block of trials from the 75 most reliable voxels from each independently defined visual regions along the human OTC (see Methods; the same results were obtained when all voxels were included from each region, see Extended Figure 3-1). These regions included topographic early visual areas V1 to V4 and higher visual object processing regions LOT and VOT (Figure 1c). LOT and VOT have been considered as the homologue of the macaque inferio-temproal (IT) cortex involved in visual object processing (Orban et al., 2004). Their responses have been shown to correlate with successful visual object detection and identification (Grill-Spector et al. 2000; Williams et al., 2007) and their lesions have been linked to visual object agnosia (Goodale et al.,1991; Farah, 2004).

The 14 CNNs we examined here included both shallower networks, such as Alexnet, VGG16 and VGG 19, and deeper networks, such as Googlenet, Inception-v3, Resnet-50 and Resnet-101 (Table 1). We also included a recurrent network, Cornet-S, that has been shown to capture the recurrent processing in macaque IT cortex with a shallower structure (Kubilius et al., 2019; Kar et al., 2019). This CNN has been recently argued to be the current best model of the primate ventral visual processing regions (Kar et al., 2019). To understand how the specific training images would impact CNN representations, besides CNNs trained with ImageNet images (Deng et al., 2009), we also examined Resnet-50 trained with stylized ImageNet images (Geirhos et al., 2019). Following a previous study (O’Connor et al., 2018), we sampled from 6 to 11 mostly pooling layers of each CNN included (see Table 1 for the specific CNN layers sampled). Pooling layers were selected because they typically mark the end of processing for a block of layers before information is passed on to the next block of layers.

### Visualizing feature coding in human OTC and CNNs

To directly visualizing how object identity and nonidentity features may be represented together in a brain region, from the z-normalized fMRI response patterns averaged over all the runs, we first calculated all pairwise Euclidean distances including all the categories shown and both values of each nonidentity feature (e.g., small and large). We then constructed a category RDM for each brain region (Kriegeskorte & Kievit, 2013, see Figure 1e). Using MDS (Shepard, 1980), we visualized this RDM by projecting the first two dimensions that captured most of the variance onto a 2D space with the distance denoting the similarity in representation among the different feature combinations (Figure 2).

A striking feature of these MDS plots was the presence or absence of the separation between the objects across the two values of each nonidentity feature. In V1, all four types of nonidentity features resulted in some separation in the representational space such that objects sharing the same value were grouped together. As information processing ascended the visual pathway, in VOT, while the separation remained visible for the two Euclidean features positions and sizes, object representations became largely overlapping for the two non-Euclidean features image stats and SF values (Figure 2).

Applying the same procedure, we also visualized how object identity and nonidentity features may be represented together in a CNN layer. Just like in early visual areas, a separation for the different values of the four nonidentity features was present in the lower layers of all 14 CNNs (see Figure 2 for results from three representative CNNs examples; see Extended Figures 2-1 to 2-4 for all the CNNs examined). In the last fully connected layer or the pooling layer before that, however, not all CNNs behaved like the higher visual regions in their encoding of position, size and image stats, and, critically, none resembled the brain in SF coding. The CNNs examined here thus do not appear to fully follow all the feature coding characteristics of the human brain, especially in higher layers.

### Quantifying the relative coding strength of identity and nonidentity features in human OTC

To quantify the relative coding strength of object identity and nonidentity features in a brain region and a CNN layer, from the z-normalized fMRI response patterns, we obtained the averaged Euclidean distance between different object categories sharing the same value of a nonidentity feature (*d_within_*) and the averaged Euclidean distance between the same object category across the two values of a nonidentity feature (*d_between_*). We then constructed an *identity dominance index* (see Methods). An identity-only representation that disregards the nonidentity feature would have a *d_between_* of “0” and an *identity dominance* of “1”. Conversely, a nonidentity-only representation that disregards identity would have a *d_within_* of “0” and an *identity dominance* of “-1”. An *identity dominance* of “0” would indicate equal representational strength of identity and nonidentity features, such that an object category is equally similar to itself in the other value of the nonidentity feature as it is to the other categories sharing the same value of the nonidentity feature. Figures 3a and 3b illustrate two scenarios of how the relative encoding strength of identity and nonidentity features may change the *identity dominance* measure between two hypothetical brain regions.

For all four types of nonidentity features examined, *identity dominance* was greater for higher than lower visual areas (the averages of LOT and VOT were all higher than those of V1 and V2, *ts* > 3.24, *ps* < .01; all reported t tests were two-tailed and corrected for multiple comparisons using the Benjamini and Hochberg method, Benjamini & Hochberg, 1995). Thus, object category encoding strength increased for all four types of nonidentity features as information ascended the visual processing pathway. Nonetheless, differences existed among them, especially between the Euclidean and non-Euclidean features (Figure 3c).

The two Euclidean features exhibited an overall similar response pattern. For position, *identity dominance* for early visual areas was very negative (the average of V1 and V2 was much lower than zero, *t(6)* = 57.04, *p* < 0.001), but increased to be close to zero in higher visual areas (the average of VOT and LOT was still less than zero, *t(6)* = 7.23, *p* = 0.005) (Figure 3c). *Identity dominance* did not vary significantly across V1 to V3 (*F(2,12)* = 3.32, p > .071), but did increase between V3 and V4, and between V4 and LOT (*ts* > 4.12, *ps* < .016), with no difference between LOT and VOT (*t(6)* = 1.54, *p* = .18). For size, *identity dominance* for early visual areas was also very negative (the average of V1 and V2 was much lower than zero, *t(6)* =16.88, *p* < 0.001), but increased to be no different from zero in higher visual areas (the average of VOT and LOT to zero, *t(6)* = 0.21, *p* = 0.84) (Figure 3c). Like position, *identity dominance* did not vary significantly from V1 to V3 (*F(2,12)* = .23, *p* > .79), but did increase between V3 and V4, and between V4 and LOT (*ts* > 3.85, *ps* < .021), with no difference between LOT and VOT (*t(6)* = 0.50, *p* = 1). These results indicated that position and size were much more prominent than objects in determining the representational space in early visual areas. The dominance of these two features over objects did not change from V1 to V3. Starting from V4, however, the strength of object coding increased until at higher visual regions LOT and VOT where it played a more or less similar role as position and size in shaping the representational space. Higher visual representations were thus never truly dominated by object identities, but rather maintained position and size information as part of the object representation.

A different pattern emerged for the two non-Euclidean features. For image stats, *identity dominance* started close to zero in early visual areas (the average of V1 and V2 was greater than zero, *paired t(5)* = 4.02, *p* = 0.014; however, V1 alone was only marginally significantly different from zero, *t(5)* = 2.22, *p* = 0.077) to significantly above zero in higher visual areas (the average of LOT and VOT was greater than zero, *t(5)* = 10.40, *p* = 0.004) (Figure 3c). *Identity dominance* also differed among V1 to V4 (*F(3,15)* = 12.94, *p* < .001) and linearly increased from V1 to V4 (with the averaged linear correlation coefficients being 0.84, and different from 0, *t(5)* = 4.92, *p* = 0.004). Pairwise tests showed that the difference between V1 and V2, and that between V4 and LOT were significant (*ts* > 3.69, *ps* < .036), with no difference observed between other adjacent region (*ts* < 2.42, *ps* > .10). Thus, while object identity and image stats were more or less equally prominent in determining object representation in early visual areas, the strength of identity coding gradually increased and dominated representation at higher levels of visual processing. For SF, although a similar overall trend of *identity dominance* going from zero in early visual areas to significantly above zero in higher visual areas was seen (*t(9)* = 0.40, *p* = 0.70, and *t(9)* = 4.79, *p* = 0.001, respectively, for the difference between zero and the averages of V1 and V2 and that of LOT and VOT), this was primarily driven by an *identity dominance* increase in LOT and VOT as *identity dominance* did not vary from V1 to V4 (no difference among V1 to V4, *F(3,27)* = 1.75, *p* = .18, the averaged linear correlation coefficient was −0.03 and no different from 0, *t(9)* = 0.39, *p* = 0.71). Thus, unlike in image stats, object identity and SF played a similar role in determining the representational space from V1 to V4; identity dominated SF representation only during higher levels of visual processing. Overall, image stats and SF appeared to be equally prominent as object identity in determining the representational space in early visual areas. As information ascends the visual processing hierarchy, however, object identity, rather than image stats or SF, took over the representational space.

To understand how the four types of nonidentity features may differ from each other, we also directly compared *identity dominance* for these four features across the four experiments using unpaired t tests (Figure 3d). In early visual regions, *identity dominance* for the average of V1 and V2 was lower for position than size (*t(12)* = 6.21, *p* < 0.001; all t tests reported here were corrected for multiple comparisons), lower for size than for both SF and image stats (*ts* > 11.05, *ps* < 0.0001), with the difference between the latter two approaching significance (*t(14)* = 2.05, *p* = 0.060). In higher visual regions, *identity dominance* for the average of LOT and VOT followed the same relative difference as in the average of V1 and V2, with all the above pairwise comparisons reaching significance (*ts* >3.06, *ps* < 0.0079).

Overall, we found that the representational strength of object identity information significantly increased over nonidentity information from lower to higher visual areas. Nevertheless, differences existed among the different nonidentity features, with identity dominating the two non-Euclidean features (image statistics and SF) but not the two Euclidean features (position and size) in higher OTC regions.

### Quantifying the relative coding strength of identity and nonidentity features in CNNs

Like the regions in OTC, for position, all CNNs tested showed very negative *identity dominance* in the early layers (Figure 3c). However, 12 of the 14 CNNs showed above zero *identity dominance* in the final layers and thus a dominance of identity over position coding not seen in the brain. For size, *identity dominance* of CNNs followed a similar trajectory as those of the brain, being very negative in the early layers and became close to zero in the final layers. Nevertheless, 13 of the 14 CNNs showed a dip in the middle layers not seen in the brain. For image stats, although, like the brain, *identity dominance* of a majority of the CNNs started close to zero in early layers and then became above zero in the final layers, their *identity dominance* trajectories differed from the brain: instead of showing a gradual increase across the layers, CNNs either showed no increase across multiple layers, or a dip below zero in the middle layers, before a rise was seen towards the end of the processing. For SF, *identity dominance* from CNNs were largely negative and none were above zero in the final layers like the brain. Overall, CNNs exhibited an over representation of identity over position and an under representation of identity over SF across low to high layers. Even for size and image stats where brain-like *identity dominance* was seen in the early and final layers, their trajectories, however, deviated from those of the brain.

As with the brain data, we also directly compared CNN *identity dominance* for the four types of nonidentity features and with those obtained from the brain (Figure 3d). While CNN lower layers exhibited a globally similar *identity dominance* profile across the four types of nonidentity features as that seen in early visual areas, the profile for the higher layers deviated from that of the higher visual regions, notably with an overrepresentation of identity over position and an underrepresentation of identity over SF (and also for image stats to some extent). Thus, while the relative coding strength of object identity and nonidentity features in lower CNN layers matched with that in early visual areas, the match between higher CNN layers and higher human visual regions were limited.

### The effect of training a CNN on original vs stylized image-net images

Although CNNs are believed to explicitly represent object shapes in the higher layers (Kriegeskorte, 2015; LeCun et al., 2015; Kubilius et al., 2016), emerging evidence suggests that CNNs may largely use local texture patches to achieve successful object classification (Ballester & de Araujo, 2016, Gatys et al., 2017) or local rather than global shape contours for object recognition (Baker et al., 2018). In a recent demonstration, CNNs were found to be poor at classifying objects defined by silhouettes and edges, and when texture and shape cues were in conflict, classifying objects according to texture rather than shape cues (Geirhos et al., 2019; see also Baker et al., 2018). However, when Resnet-50 was trained with stylized ImageNet images in which the original texture of every single image was replaced with the style of a randomly chosen painting, object classification performance significantly improved, relied more on shape than texture cues, and became more robust to noise and image distortions (Geirhos et al., 2019). It thus appears that a suitable data set may overcome the texture bias in standard CNNs and allow them to utilize more shape cues.

Here we tested if the relative coding strength of object identity and nonidentity features in a CNN may become more brain-like when the CNN was trained with stylized ImageNet images. To do so, we examined the representations formed in Resnet-50 pretrained with three different procedures (Geirhos et al., 2019): trained only with the stylized ImageNet Images (RN50-SIN), trained with both the original and the stylized ImageNet Images (RN50-SININ), and trained with both sets of images and then fine-tuned with the stylized ImageNet images (RN50-SININ-IN). For comparison, we also included Resnet-50 trained with the original ImageNet images (RN50-IN) that we tested before.

Despite minor differences, *object dominance* measures were remarkably similar whether or not Resnet-50 was trained with the original or the stylized ImageNet images: all were still substantially different from the human ventral regions (Figure 3e). Thus, despite the improvement in object classification performance with the inclusion of the stylized images, the relative coding strength of object identity and nonidentity features in Resnet-50 did not appear to change. The incorporation of stylized ImageNet images likely forced Resnet-50 to use long-range structures rather than local texture structures, but without fundamentally changing how images were computed. The inability of Resnet-50 to exhibit brain-like *object dominance* suggests that such a difference could not be overcome by this type of training.

## DISCUSSION

Despite the usefulness of object identity and nonidentity features in vision and their joint coding in visual processing in the primate brain, they have so far been studied relatively independently. Here we documented the relative coding strength of object identity and nonidentity features within a brain region and tracked how it changed over the course of processing along the human ventral visual pathway. We compared object identity with nonidentity features, which included both Euclidean features (position and size) and non-Euclidean features (basic image statistics and the SF content of an image). We additionally compared responses from the human brain with those from 14 CNNs pretrained for object recognition.

Our measure of the relative coding strength of object identity and nonidentity features depended on the variation we introduced in each feature. For example, the relative coding strength of two similar object identities over two dissimilar object positions could be very different from that of two different object identities over two similar object positions. Because similarity within a feature dimension changes across brain regions (e.g., similar objects in one region may become dissimilar in another region), it would not have been possible to equate feature variations for all the brain regions examined. Thus we have chosen what we believed to be reasonable variations for each feature, including sampling a wide range of real-world object categories, choosing two positions and two sizes that were as different as possible but still allowed each object at a given position or size to be visible, and choosing two SF ranges that were far apart from each other. The goal here was therefore not to measure the absolute feature coding bias in a brain region or a CNN layer, but rather, for a set of reasonably chosen features, how the feature coding bias may change across visual areas and CNN layers to inform us how object identity and nonidentity features may be represented together.

Overall, we found that the encoding of object identity significantly increased over all four types of nonidentity features from lower to higher visual areas. Meanwhile, differences existed among these different nonidentity features, with identity dominating the non-Euclidean features (image statistics and SF) but not the Euclidean features (position and size) in higher OTC regions. Specifically, position and size were much more prominent than objects in determining the representational space in early visual areas. The dominance of these two features over objects did not change from V1 to V3. Starting from V4, the strength of object coding increased until at higher visual regions LOT and VOT where it played a more or less similar role as position and size in shaping the representational space. Higher visual representations were thus never truly dominated by object identities, but rather maintaining position and size information as part of the object representation. This is consistent with the existence of topographic maps in higher visual regions (Brewer et al. 2005) and the robust representation of position information in monkey IT (DiCarlo & Maunsell, 2003; Hung et al., 2004; Hong et al., 2016) and in human VOT and LOT (Schwarzlose et al., 2008; Kravitz et al., 2008 & 2010; Carlson et al., 2011; Cichy et al., 2011 & 2013). Meanwhile, image stats and SF were equally prominent as identity in determining the representational space in early visual areas. As information ascends the visual processing hierarchy, however, identity dominated image stats and SF in the representational space. While the dominance of identity over image stats increased steadily from V1 to LOT/VOT, the dominance of identity over SF remained relatively stable from V1 to V4 and then increased significantly in LOT/VOT. Together, our study documented for the first time the relatively coding strength of object identity and nonidentity features in different human visual processing regions and its evolution of along the ventral processing pathway.

Why would identity dominate the two non-Euclidean but not the two Euclidean features examined at higher levels of ventral visual processing? One possibility could be that Euclidean features are more essential in our direct interaction with the objects, such as in reaching and grasping. The non-Euclidean features, on the other hand, may be discarded once object identity and other non-identity information is recovered. For example, once we identity an object and its position and size in a foggy viewing condition, information about how foggy it is may no longer be useful in guiding our further interaction with the object. Further studies are needed to verify this speculation.

Compared to regions in human OTC, the 14 CNNs we examined exhibited an over representation of identity over position and an under representation of identity over SF from lower to higher layers. Even for size and image stats where brain-like responses were seen in the early and final layers, their trajectories, however, deviated from those of the brain. Direct comparison of the encoding of the four types of nonidentity features with respect to object identity revealed that while CNN lower layers exhibited a globally similar response profile across the four types of nonidentity features as that seen in early visual areas, the profile for the higher layers deviated from that of the higher visual regions. Thus, while the relative coding strength of object identity and nonidentity features in lower CNN layers of CNNs matched with that of early human visual areas, the match between higher CNN layers and higher human visual regions were limited. Similar results were obtained for the different CNNs tested and for a CNN trained with stylized object images that emphasized shape representation.

To increase SNR, using a block design, we averaged responses from multiple exemplars from the same category. We thus examined object category responses rather than responses to each induvial exemplar. Previous research has shown similar category and exemplar response profiles in macaque IT and human lateral occipital cortex with more robust responses by categories than individual exemplars due to an increase in SNR (Hung et al., 2005; Cichy et al., 2011). In a recent study, Rajalingham, et al. (2018) found better behavior-CNN correspondence at the category but not at the individual exemplar level. Thus, comparing responses averaged overall multiple exemplars at the category level, rather than at the exemplar level, should have increased our chance of finding a close brain-CNN correspondence if it existed. Yet, such a close correspondence was only found at the lower, but not at higher, levels of visual processing. This result echoes our finding in another study in which we showed that the object representational structures formed in lower CNN layers could fully capture those formed in lower human visual processing regions but that higher CNN layers failed to do so for higher human visual processing regions (Xu & Vaziri-Pashkam, 2020).

We included in this study both shallow and very deep CNNs. Deeper CNNs in general exhibit better object recognition performance (as evident from the ImageNet challenge results, see Russakovsky et al., 2015), and can partially approximate the recurrent processing in ventral visual regions (Kar et al., 2019). The recurrent CNN we examined here, Cornet-S, explicitly models recurrent processing in ventral visual areas (Kar et al., 2019) and is considered by some as the current best model of the primate ventral visual regions (Kubilius et al., 2019). Yet we observed similar performance between shallow and deep CNNs (e.g., Alexnet vs Googlenet), and the recurrent CNN did not perform better than the other CNNs.

The present results join other studies that have shown differences between the CNNs and brain, such as the kind of features used in object recognition (Ballester & de Araujo, 2016, Ulman et al., 2016; Gatys et al., 2017; Baker et al., 2018; Geirhos et al., 2019), disagreement in representational structure between CNNs and brain/behavior (Kheradpisheh et al., 2016; Karimi-Rouzbahani et al., 2017; Rajalingham, et al., 2018; Xu & Varziri-Pashkam, 2020), the inability of CNN to explain more than 55% of the variance of macaque V4 and IT neurons (Cadieu et al., 2014; Yamins et al., 2014; Kar et al. 2019; Bao et al., 2020), and how the two systems handle adversarial images (see Serre, 2019). Here by using a novel measure of documenting the relative coding strength of object identity and nonidentity features, we showed that although there was a close brain-CNN correspondence at lower levels of visual processing, such a correspondence was limited at higher levels of visual processing.

To conclude, by documenting the relative coding strength of object identity and nonidentity features in different human visual processing regions and its evolution of along the ventral processing pathway, we showed overall an increase of identity and a decrease of nonidentity information representation during the course of visual processing in the human brain. While the representation of identity dominated those of the non-Euclidean features in higher levels of visual processing, this was not the case for the Euclidean features examined. Our examination of 14 CNNs further revealed that, while feature coding in lower CNN layers matched with that of human early visual areas, the match between higher CNN layers and higher human visual regions were limited.

## Acknowledgement

We thank Martin Schrimpf for help implementing CORnet-S, JohnMark Tayler for extracting the features from the three Resnet-50 models trained with the stylized images, and Thomas O’Connell, Brian Scholl, JohnMark Taylor and Nick Turk-Brown for helpful discussions and feedback on the results. This research was supported by National Institute of Health Grants (1R01EY030854 and 1R01EY022355) to Y.X.

---

**Extended Figure 2-1.**
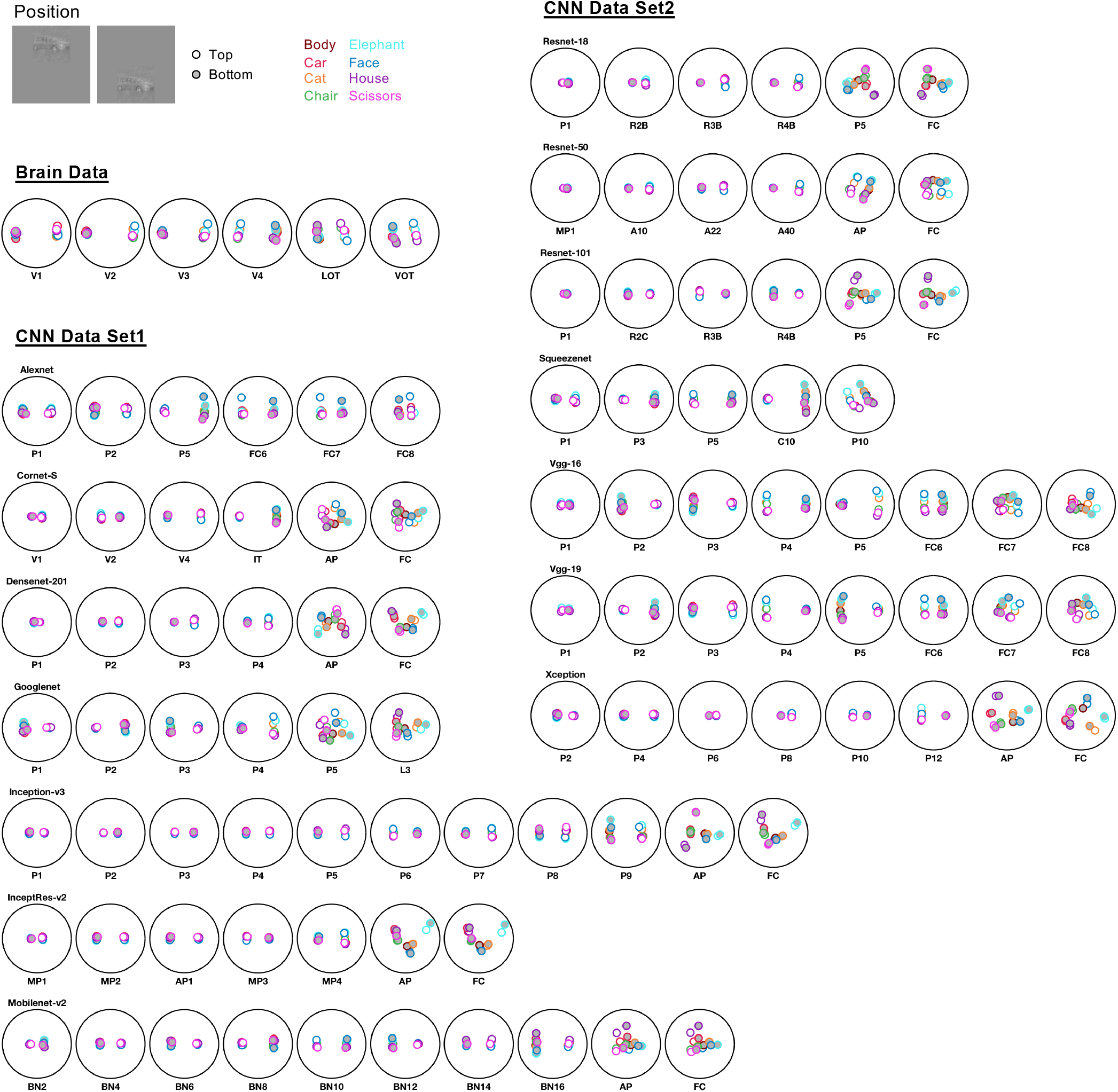
Visualizing identity and position coding in human OTC and all the CNNs examined.

**Extended Figure 2-2.**
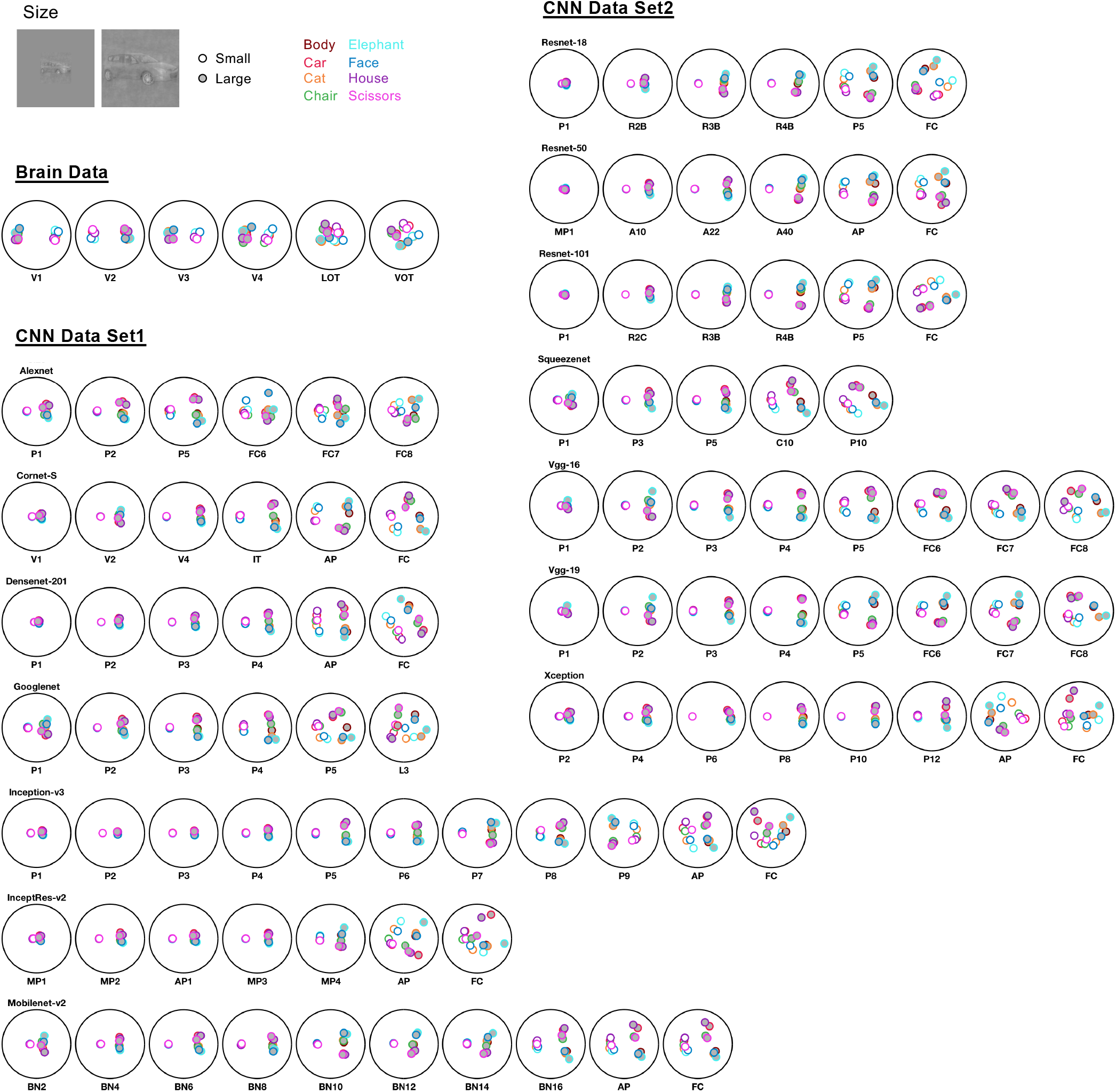
Visualizing identity and size coding in human OTC and all the CNNs examined.

**Extended Figure 2-3.**
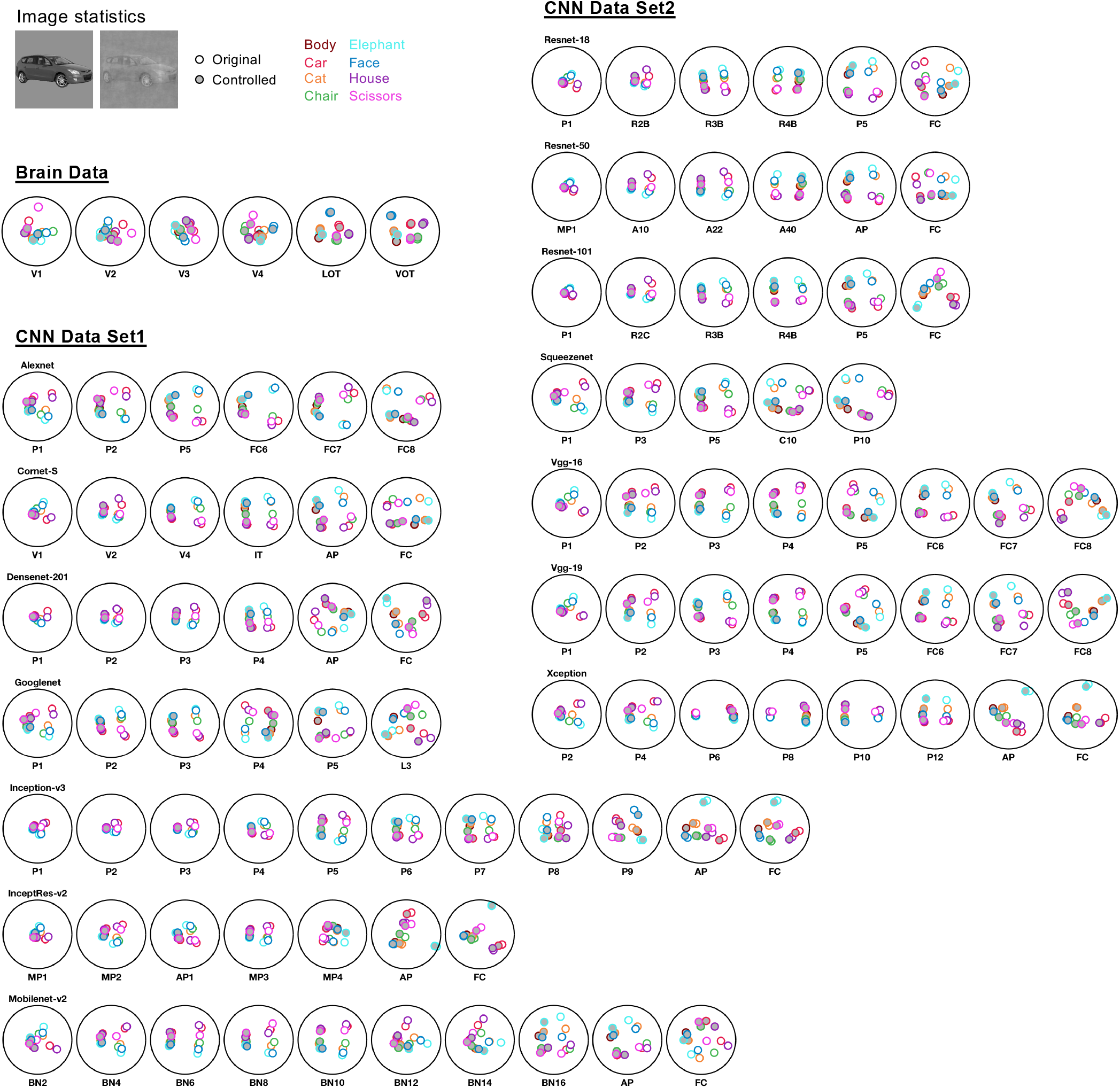
Visualizing identity and image stats coding in human OTC and all the CNNs examined.

**Extended Figure 2-4.**
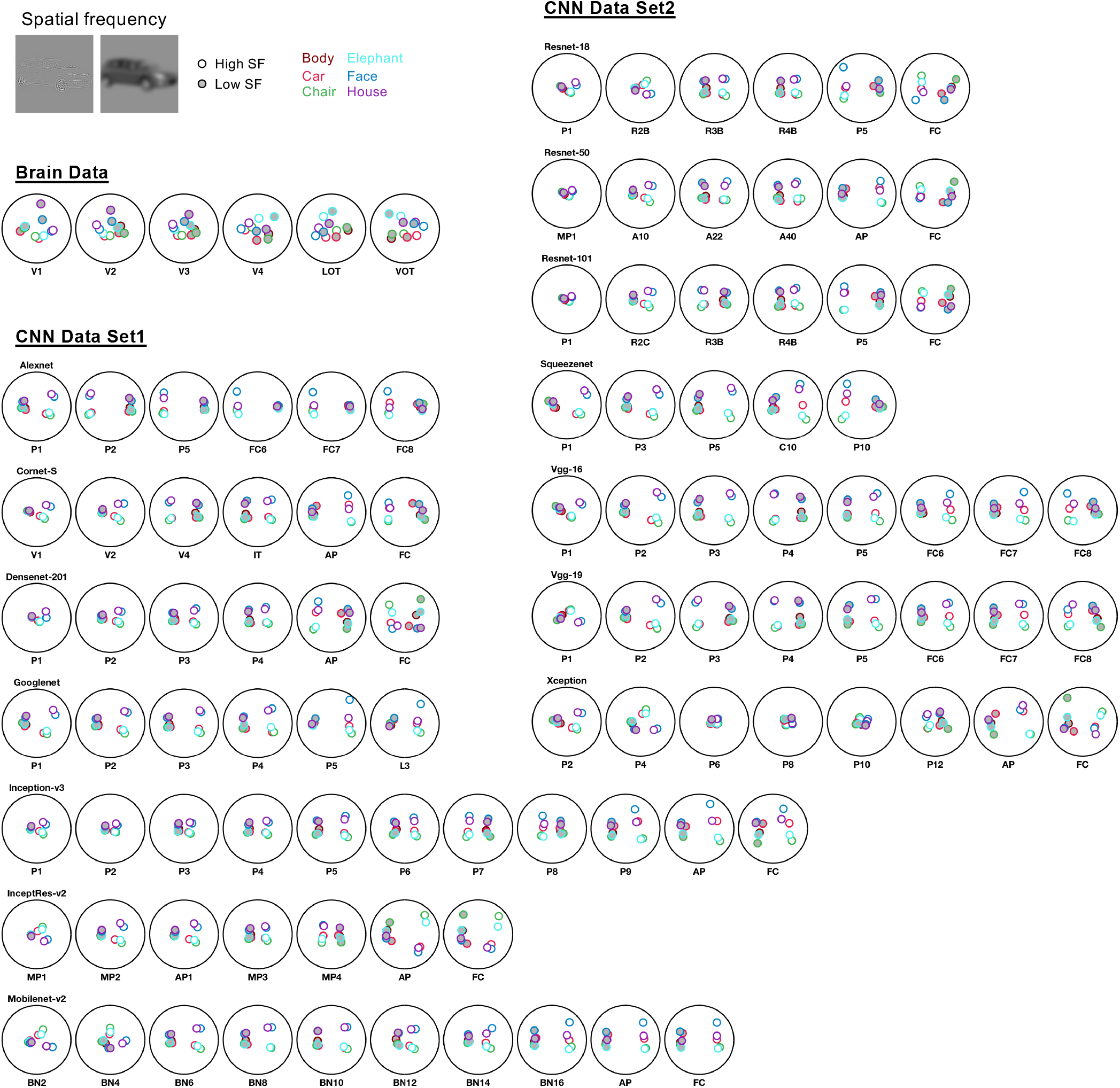
Visualizing identity and SF coding in human OTC and all the CNNs examined.

**Extended Figure 3-1.**
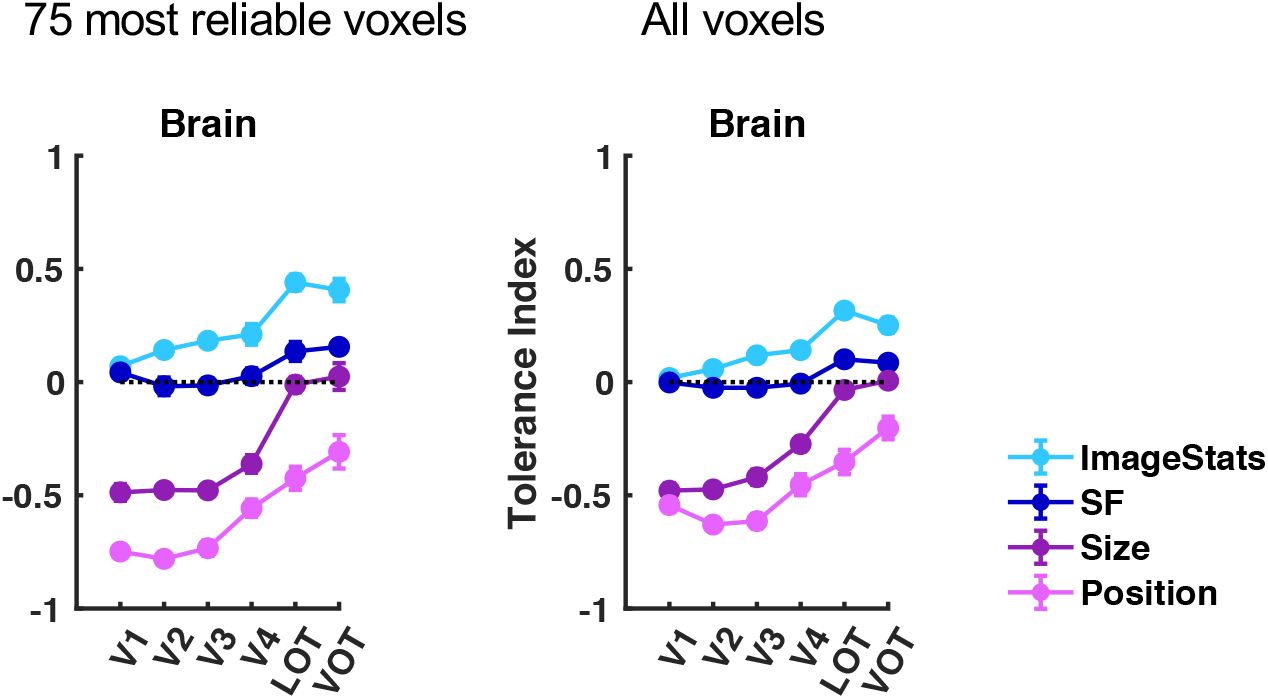
*Identity dominance* from human OTC including the 75 most reliable voxels (left) and all the voxel (right) from each brain region examined. Similar response patterns were seen in both.

## References

Baker N, Lu H, Erlikhman G, Kellman PJ (2018) Deep convolutional networks do not classify based on global object shape. PLOS Comput Biol 14:e1006613.

Ballester, P, de Araújo RM (2016) On the Performance of GoogLeNet and AlexNet Applied to Sketches. In AAAI (pp. 1124–1128).

Bao P, She L, McGill M, Tsao DY (2020) A map of object space in primate inferotemporal cortex. Nature https://doi.org/10.1038/s41586-020-2350-5.

Benjamini Y, Hochberg Y (1995) Controlling the False Discovery Rate - a Practical and Powerful Approach to Multiple Testing. J Roy Stat Soc B Met 57:289–300.

Bettencourt KC, Xu Y (2016) Understanding location- and feature-based processing along the human intraparietal sulcus. J Neurophysiol 116:1488–97.

Brewer AA, Liu J, Wade AR, Wandell BA (2005) Visual field maps and stimulus selectivity in human ventral occipital cortex. Nat Neurosci 8:1102–1109.

Cadieu CF, Hong H, Yamins DLK, Pinto N, Ardila D, et al. (2014) Deep neural networks rival the representation of primate IT cortex for core visual object recognition. PLOS Comput Biol 10:e1003963.

Carlson T, Hogendoorn H, Fonteijn H, Verstaten FAJ (2011) Spatial coding and invariance in object-selective cortex. Cortex 47:14–22.

Cichy RM, Chen Y, Haynes JD (2011) Encoding the identity and location of objects in human LOC. Neuroimage 54:2297–2307.

Cichy RM, Sterzer P, Heinzle J, Elliott LT, Ramirez F, Haynes JD (2013) Probing principles of large-scale object representation: Category Preference and location encoding. Hum Brain Mapp 34:1636–1651.

Cichy RM, Khosla A, Pantazis D, Torralba A, Oliva A (2016) Comparison of deep neural networks to spatiotemporal cortical dynamics of human visual object recognition reveals hierarchical correspondence. Sci Rep 6:27755.

Cichy RM, Kaiser D (2019) Deep neural networks as scientific models. Trends Cogn Sci 23:305–317.

Dale AM, Fischl B, Sereno MI (1999) Cortical surface-based analysis. I. Segmentation and surface reconstruction. Neuroimage 9:179–194.

Deng J, Dong W, Socher R, Li LJ, Li K, Fei-Fei L (2009) ImageNet: A largescale hierarchical image database. In IEEE conference on computer vision and pattern recognition. CVPR (pp. 248–255).

DiCarlo JJ, Cox DD (2007) Untangling invariant object recognition. Trends Cogn Sci 11: 333–341.

DiCarlo JJ, Zoccolan D, Rust RC (2012) How does the brain solve visual object recognition? Neuron 73:415–434.

DiCarlo JJ, Manusell JHR (2003) Anterior inferotemporal neurons of monkeys engaged in object recognition can be highly sensitive to object retinal position. J Neurophysiol 89:3264–3278.

Eickenberg M, Gramfort A, Varoquaux G, Thirion B (2017) Seeing it all: Convolutional network layers map the function of the human visual system. NeuroImage 152:184–94.

Farah MJ (2004) Visual agnosia. Cambridge, Mass.: MIT Press.

Gatys LA, Ecker AS, Bethge M (2017) Texture and art with deep neural networks. Curr Opin Neurobiol 46:178–86.

Geirhos R, Rubisch P, Michaelis C, Bethge M, Wichmann FA, Bren-del W (2019) ImageNet-trained CNNs are biased towards texture; increasing shape bias improves accuracy and robustness. In International Conference on Learning Representations.

Goodale MA, Milner AD, Jakobson LS, Carey DP (1991) A neurological dissociation between perceiving objects and grasping them. Nature 349:154–156.

Grill-Spector K, Kushnir T, Hendler T, Malach R (2000) The dynamics of object-selective activation correlate with recognition performance in humans. Nat Neurosci 3:837–843.

Grill-Spector K, Kushnir T, Edelman S, Itzchak Y, Malach R (1998) Cue-invariant activation in object-related areas of the human occipital lobe. Neuron 21:191–202.

Güçlü U, van Gerven MAJ (2017) Increasingly complex representations of natural movies across the dorsal stream are shared between subjects. NeuroImage 145:329–36.

Hong H, Yamins DLK, Majaj NJ, DiCarlo JJ (2016) Explicit information for category-orthogonal object properties increases along the ventral stream. Nat Neurosci 19: 613–622.

Hung CP, Kreiman G, Poggio T, DiCarlo JJ (2005) Fast readout of object identity from macaque inferior temporal cortex. Science 310:863–866.

Kar K, Kubilius J, Schmidt K, Issa EB, DiCarlo JJ (2019) Evidence that recurrent circuits are critical to the ventral stream’s execution of core object recognition behavior. Nat Neurosci 22:974–983.

Karimi-Rouzbahani H, Bagheri N, Ebrahimpour R (2017) Invariant object recognition is a personalized selection of invariant features in humans, not simply explained by hierarchical feedforward vision models. Sci Rep 7: 14402.

Khaligh-Razavi S-M, Kriegeskorte N (2014) Deep supervised, but not unsupervised, models may explain IT cortical representation. PLOS Comput Biol 10:e1003915.

Kheradpisheh SR, Ghodrati M, Ganjtabesh M, Masquelier T (2016) Deep networks can resemble human feed-forward vision in invariant object recognition. Sci Rep 6:32672.

Kourtzi Z, Kanwisher N (2000) Cortical regions involved in perceiving object shape. J Neurosci 20:3310–3318.

Kravitz DJ, Vinson LD, Baker CI (2008) How position dependent is visual object recognition? Trends Cogn Sci 12:114–122.

Kravitz DJ, Kriegeskorte N, Baker CI (2010) High-level visual object representations are constrained by position. Cereb Cortex 20:2916–2925.

Kriegeskorte N (2015) Deep neural networks: a new framework for modeling biological vision and brain information processing. Annu Rev Vis Sci 1:417–46.

Kriegeskorte N, Kievit RA (2013) Representational geometry: integrating cognition, computation, and the brain. Trends Cogn Sci 17:401–412.

Kubilius J, Bracci S, Op de Beeck HP (2016) Deep neural networks as a computational model for human shape sensitivity. PLOS Comput Biol 12:e1004896.

Kubilius J, Schrimpf M, Hong H, et al. (2019) Brain-like object recognition with high-performing shallow recurrent ANNs. In: Neural Information Processing Systems. Vancouver, British Columbia, Canada.

LeCun Y, Bengio Y, Hinton G (2015) Deep learning. Nature 521:436–444.

Malach R, Reppas JB, Benson RR, Kwong KK, Jiang H, Kennedy WA, Ledden PJ, Brady TJ, Rosen BR, Tootell RB (1995) Object-related activity revealed by functional magnetic resonance imaging in human occipital cortex. Proc Nati Acad Sci USA 92:8135–8139.

O’Connell TP, Chun MM (2018) Predicting eye movement patterns from fMRI responses to natural scenes. Nat. Commun 9, 5159.

Orban GA, Van Essen D, Vanduffel W (2004) Comparative mapping of higher visual areas in monkeys and humans. Trends Cogn Sci 8:315–324.

Rajalingham R, Issa EB, Bashivan P, Kar K, Schmidt K, DiCarlo JJ (2018) Large-scale, high-resolution comparison of the core visual object recognition behavior of humans, monkeys, and state-of-the-art deep artificial neural networks. J Neurosci 38:7255–69.

Russakovsky O, Deng J, Su H, Krause J, Satheesh S, et al. (2015) ImageNet Large Scale Visual Recognition Challenge. Int J Comput Vis 115:211–52.

Schwarzlose RF, Swisher JD, Dang S, Kanwisher N (2008) The distribution of category and location information across object-selective regions in human visual cortex. Proc Nati Acad Sci USA 105: 4447–4452.

Serre T (2019) Deep learning: The good, the bad, and the ugly. Annu. Rev. Vis. Sci. 5:21.1–21.28.

Shepard RN (1980) Multidimensional scaling, tree-fitting, and clustering. Science 210:390–398.

Swisher JD, Halko MA, Merabet LB, McMains SA, Somers DC (2007) Visual Topography of Human Intraparietal Sulcus. J Neurosci 27:5326–5337.

Tahan L, Konkle T (2020) Reliability-based voxel selection. Neuroimage 207:116350.

Ullman S, Assif L, Fetaya E, Harari D (2016) Atoms of recognition in human and computer vision. Proc Nati Acad Sci USA 113:2744–49.

Vaziri-Pashkam M, Xu Y (2019) An information-driven two-pathway characterization of occipito-temporal and posterior parietal visual object representations. Cereb Cortex 29:2034–2050.

Vaziri-Pashkam M, Taylor J, Xu Y (2019) Spatial frequency tolerant visual object representations in the human ventral and dorsal visual processing pathways. J Cogn Neurosci 31:49–63.

Willenbockel V, Sadr J, Fiset D, Horne GO, Gosselin F, Tanaka JW (2010) Controlling low-level image properties: The SHINE toolbox. Behavior Research Methods 42:671–684.

Williams MA, Dang S, Kanwisher NG (2007) Only some spatial patterns of fMRI response are read out in task performance. Nat Neurosci 10:685–686.

Xu Y, Vaziri-Pashkam M (2020) Limited correspondence in visual representation between the human brain and convolutional neural networks. bioRxiv.

Yamins DLK, DiCarlo JJ (2016) Using goal-driven deep learning models to understand sensory cortex. Nat Neurosci 19:356–65.

Yamins DLK, Hong H, Cadieu CF, Solomon EA, Seibert D, DiCarlo JJ (2014) Performance-optimized hierarchical models predict neural responses in higher visual cortex. Proc Nati Acad Sci USA 111:8619–24.

